# *Copb2* is essential for embryogenesis and hypomorphic mutations cause human microcephaly

**DOI:** 10.1101/135236

**Authors:** Andrew DiStasio, Ashley Driver, Kristen Sund, Milene Donlin, Ranjith M. Muraleedharan, Shabnam Pooya, Beth Kline-Fath, Kenneth M. Kaufman, Cynthia A. Prows, Elizabeth Schorry, Biplab DasGupta, Rolf W. Stottmann

## Abstract

Primary microcephaly is a congenital brain malformation characterized by a head circumference less than three standard deviations below the mean for age and sex and results in moderate to severe mental deficiencies and decreased lifespan. We recently studied two children with primary microcephaly in an otherwise unaffected family. Exome sequencing identified an autosomal recessive mutation leading to an amino acid substitution in a WD40 domain of the highly conserved *Coatomer Protein Complex, Subunit Beta 2 (COPB2)*. To study the role of *Copb2* in neural development, we utilized genome editing technology to generate an allelic series in the mouse. Two independent null alleles revealed that *Copb2* is essential for early stages of embryogenesis. Mice homozygous for the patient variant (*Copb2^R^254^C/R^254^C^*) appear to have a grossly normal phenotype, likely due to differences in corticogenesis between the two species. Strikingly, mice heterozygous for the patient mutation and a null allele (*Copb2^R^254^C/Znf^*) show a severe perinatal phenotype including low neonatal weight, significantly increased apoptosis in the brain, and death within the first week of life. Immunostaining of the *Copb2^R^254^C/Znf^* brain revealed a reduction in layer V (CTIP2^+^) neurons, while the overall cell density of the cortex is unchanged. Moreover, disruption of *Copb2* in mouse neurospheres resulted in reduced proliferation. These results identify a general requirement for *COPB2* in embryogenesis and a specific role in corticogenesis. We further demonstrate the utility of CRISPR-Cas9 generated mouse models in the study of potential pathogenicity of variants of potential clinical interest.

## INTRODUCTION

Autosomal recessive primary microcephaly is a rare congenital brain defect resulting in a reduction of occiptofrontal head circumference by at least 3 standard deviations. Patients with microcephaly are typically afflicted with varying degrees of intellectual disability and can either be part of a syndromic condition or an isolated malformation (1, 2). To date, mutations in at least twelve genes have been identified in patients with primary microcephaly. The roles of these genes include proper orientation of the mitotic spindle, the duplication, formation or function of the centrosome, DNA-damage response, transcriptional regulation and vesicle trafficking. Most of these mutated genes ultimately play roles in neural progenitor proliferation during the intricate development and organization of the cerebral cortex (3-11).

While there has been significant progress identifying genetic causes of primary microcephaly, the genetics of neurodevelopmental disorders remains incompletely understood (12). In these cases, exome sequencing has become an efficient tool for the identification of rare disease variants. We performed exome sequencing in a pedigree with two affected siblings with primary microcephaly and identified a homozygous recessive variant in the coding region of *Coatomer Protein Complex Subunit Beta 2 (COPB2).*

*COPB2* encodes β’-COP, a subunit of the Golgi Coatomer complex (COPI), which is necessary for retrograde trafficking from the Golgi to the Endoplasmic Reticulum (ER) (13-16). In addition to its structural role in the COPI complex, β’-COP has been shown to directly interact with cargo (17, 18) and with regulators of intracellular trafficking. In order to test our hypothesis that the microcephaly and other congenital malformations seen in the affected patients are due to the missense mutation in the coding region of *COPB2*, we used genome editing technology to generate an allelic series in the mouse. Our results indicate that complete loss of *Copb2* is incompatible with embryonic survival to organogenesis stage of embryonic development, and that compound heterozygotes for a null allele and the missense allele of *Copb2* show both brain and growth defects in mice. Thus, we have identified a novel causal gene for primary microcephaly and a requirement for this gene in early mammalian development.

## RESULTS

### Homozygous recessive mutation in COPB2 identified in two patients with primary microcephaly

Two affected children presented in our hospital with abnormal head size, severe developmental delay, failure to thrive, cortical blindness and spasticity. Both parents and a sibling were healthy with no features of microcephaly (Fig 1A). Magnetic resonance imaging confirmed microcephaly with a simplified gyral pattern, thin corpus callosum, slight dilation of the lateral, third and fourth ventricles, enlarged extra-axial space and delayed myelination (Fig 1B-E). Occipital head circumference was well below the 3^rd^ percentile at all ages for both affected children (Table 1). The low body weight and proportional reduction in head size became more severe in each child as they aged (Fig 1 F-I). In contrast to these measurements, the height of both children was well within the range of normal growth. To further define the microcephaly phenotype based on clinical classifications (2) and to determine whether the clinical phenotype could differentiate between a role for proliferation and/or atrophy in the underlying mechanism, we performed a detailed analysis of the brain imaging studies. We measured the extra-axial fluid spaces in the area of the cisterna magna and sylvian fissures and sizes for the atrium of the lateral, third and fourth ventricles over development of the affected children. This analysis showed a progressive increase in brain ventricle size and minimal change in extra-axial space dimensions (Table 2) suggesting progressive central brain volume loss.

**Figure 1.**
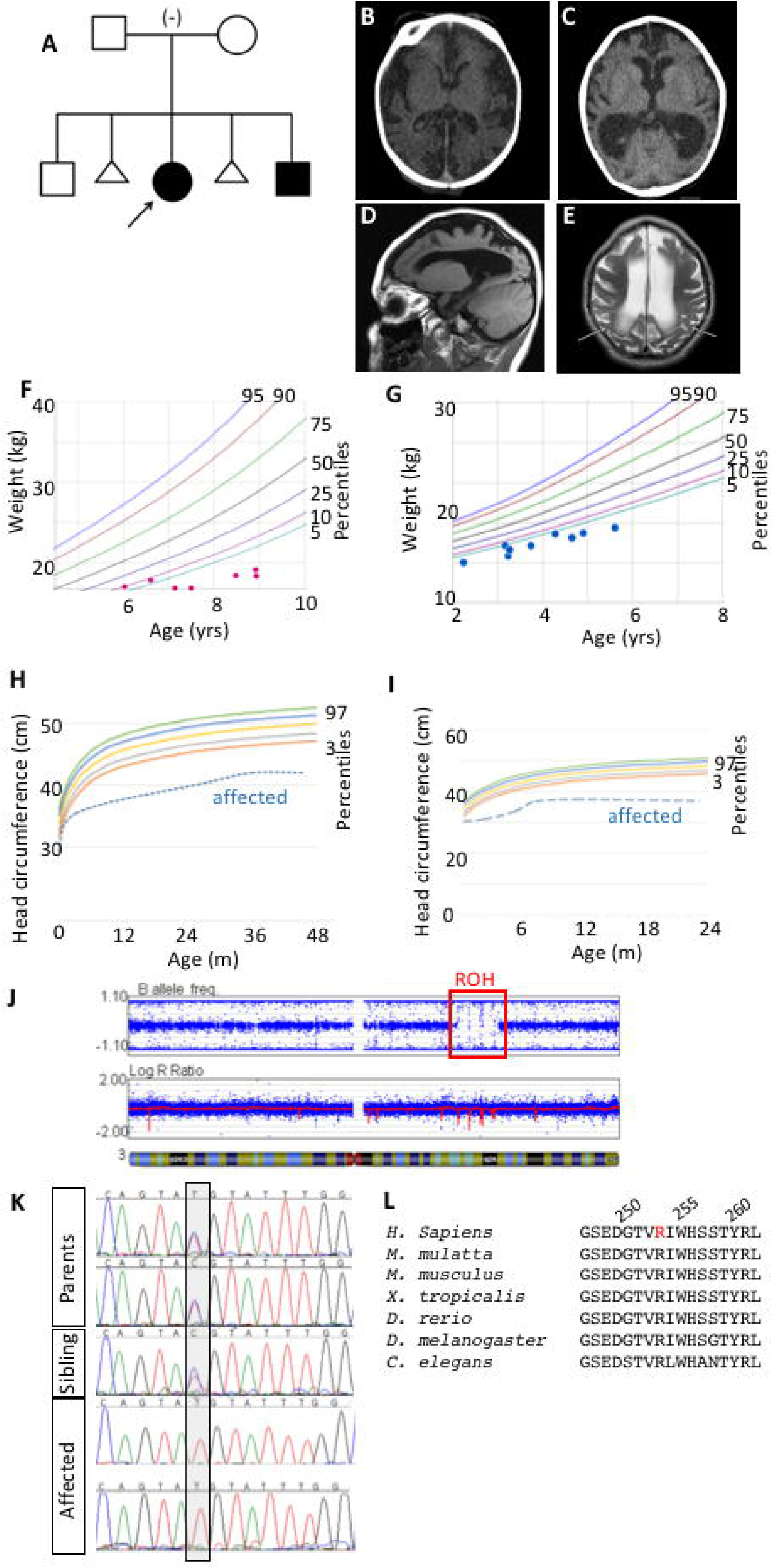
Microcephaly in two patients with a single region of homozygosity on chromosome 3. (A) Pedigree of family with autosomal recessive primary microcephaly in two affected children with unaffected parents and sibling. (Proband is indicated with arrow) (B, C) Computed tomography (CT) scan of the proband (B) and the affected sibling (C) revealed decreased sulci and enlarged extra axial fluid spaces and ventricles. Sagittal T1 weighted MR image of the proband at age 4.5 (D) depicts microcephaly with a simplified gyral pattern, dilation of the lateral ventricle, and enlarged extra axial space. Axial T2 weighted image of the proband (E) again demonstrates simplified gyral pattern and enlarged lateral ventricles and extra axial spaces with increased signal in the white matter (arrows) consistent with delayed myelination. (F-I) The proband (F,H) and affected sibling (G,I) are consistently below 5^th^ percentile in weight and below the 3^rd^ percentile for head circumference. (J) SNP microarray analysis identified a single large region of homozygosity (ROH, red box) was identified in both affected siblings at 3q22.1q24. Parents denied consanguinity and all other ROH were less than 2.6 Mb in size. (K) Sanger Sequencing confirms exome results that affected children are homozygous for the n.760C>T sequence variant while both parents and the unaffected sibling are heterozygous. (L) The mutation results in a p.R254C coding change in a highly conserved region of the protein (mutated residue in red, human amino acid positions 250,255,260 indicated).

**Table 1.**
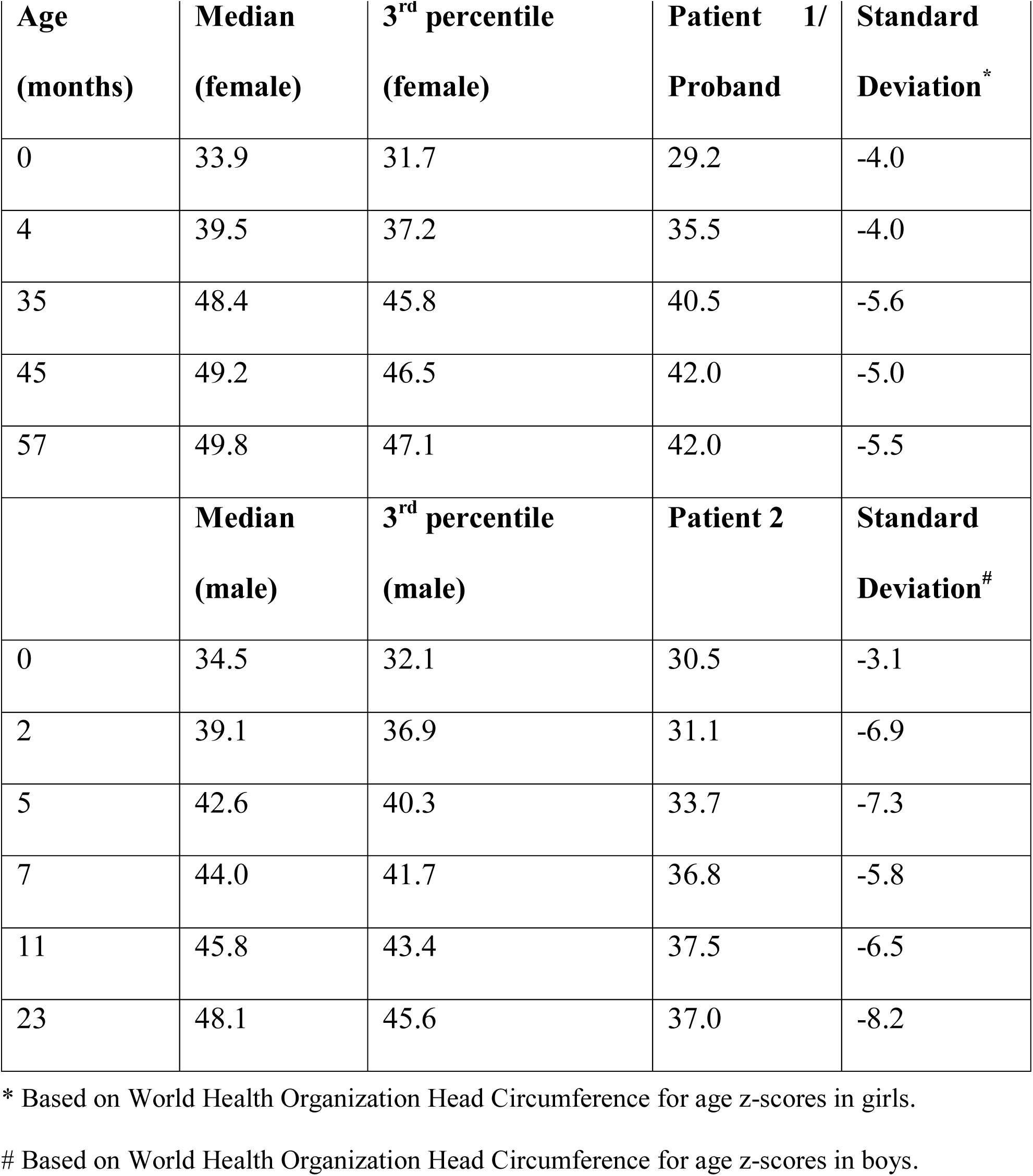
Occipital head circumference measurement (cm) and standard deviation from mean WHO head circumference over time.

| **Age**<br>**(months)** | **Median**<br>**(female)** | **3<sup>rd</sup> percentile**<br>**(female)** | **Patient 1/**<br>**Proband** | **Standard**<br>**Deviation \*** |
| --- | --- | --- | --- | --- |
| 0 | 33.9 | 31.7 | 29.2 | -4.0 |
| 4 | 39.5 | 37.2 | 35.5 | -4.0 |
| 35 | 48.4 | 45.8 | 40.5 | -5.6 |
| 45 | 49.2 | 46.5 | 42.0 | -5.0 |
| 57 | 49.8 | 47.1 | 42.0 | -5.5 |
|  | **Median**<br>**(male)** | **3<sup>rd</sup> percentile**<br>**(male)** | **Patient 2** | **Standard**<br>**Deviation<sup>#</sup>** |
| 0 | 34.5 | 32.1 | 30.5 | -3.1 |
| 2 | 39.1 | 36.9 | 31.1 | -6.9 |
| 5 | 42.6 | 40.3 | 33.7 | -7.3 |
| 7 | 44.0 | 41.7 | 36.8 | -5.8 |
| 11 | 45.8 | 43.4 | 37.5 | -6.5 |
| 23 | 48.1 | 45.6 | 37.0 | -8.2 |

\* Based on World Health Organization Head Circumference for age z-scores in girls.

**Table 2.** Measurement of extra axial space (mm) and brain ventricles (mm) demonstrate progressive enlargement of ventricles over time with minimal progressive change in the extra axial space.

| Age (months) | Extra Axial |  |  | Left Lateral | Right Lateral | Third | Forth Ventricle |  |
| --- | --- | --- | --- | --- | --- | --- | --- | --- |
|  | CM | SF-L | SF-R | Ventricle (T) * | Ventricle (T)* | Ventricle (T)* | AP | T |
| 0 | 11.9 | 13.7 | 13.9 | 19.8 | 18.8 | 6.7 | 7.3 | 7.9 |
| 13 | 11.6 | 14.0 | 14.9 | 22.7 | 23.5 | 9.4 | 9.4 | 10.7 |
| 50 | 11.6 | 15.0 | 16.4 | 28.6 | 25.4 | 10.6 | 12.0 | 15.6 |
| Age (months) | Extra Axial |  |  | Left Lateral | Right Lateral | Third | Forth Ventricle |  |
|  | CM | SF-L | SF-R | Ventricle | Ventricle | Ventricle | AP | T |
| 0 | 8.2 | 15.0 | 14.6 | 13.8 | 7.4 | 4.3 | 5.2 | 8.2 |
| 7 | 7.8 | 17.3 | 13.4 | 15.6 | 7.4 | 7.4 | 6.3 | 12.1 |
| 39 | 9.4 | 17.2 | 16.7 | 15.2 | 9.2 | 8.6 | 7.0 | 15.6 |

\* In a large study of cerebral ventricle size in children (53), the greatest transverse length of the lateral ventricle was 10 mm and the third ventricle was 5 mm.
CM= Cisterna Magna
SF-L= Left Sylvian Fissure
SF-R= Right Sylvian Fissure
AP= anterior-posterior
T= transverse

To determine the genetic origin of this syndrome, an initial series of microcephaly genes (*ASPM, CK5RAP2, CENP, MCPH1, STIL*) was sequenced in the proband with negative results. The affected siblings had a SNP microarray analysis performed using the Illumina Infinium Assay to assess copy number variation (CNV) and regions of homozygosity (ROH). Subsequent analysis did not identify any pathogenic CNV’s but did find a single large region of homozygosity (ROH) in both the proband and her affected brother, despite parental report of non-consanguinity (Fig 1J). The 16.8 Mb ROH was identified at 3q22.1q24 and contained approximately 90 Refseq genes.

Whole exome sequencing (Illumina Hi Seq 2000) was performed for the two affected children and their parents after obtaining informed consent according to our Cincinnati Children’s Hospital Medical Center (CCHMC) IRB-approved protocol. Alignment and variant detection was performed using the Broad Institute’s web-based Genome Analysis Tookit (GATK; Genome Reference Consortium Build 37) (19). Quality control and data filtering were performed on VCF files in Golden Helix’s SNP and Variation Suite. Non-synonymous coding variants were compared to three control databases, including NHLBI’s ESP6500 exome data (20), the 1000 genomes project (21), and an internal CCHMC control cohort (22). The initial bioinformatics analysis identified 126,752 variants (Table 3). After filtering for quality control using filters previously described (22), coding variants, non-synonymous variants, and minor allele frequencies, we identified one single mutation which followed a homozygous recessive inheritance pattern. The identified variant was compared to known disease genes in the OMIM and Human Gene Mutation (HGMD) (23) databases, and to reported variants in dbSNP (24) and the Exome Aggregation Consortium (ExAC; (25). There is one reported heterozygous variant in this position in the Exac database coding for a different missense variant (p.R253H). The variant was also analyzed using Interactive Biosoftware’s Alamut v2.2 to determine location of mutation within a protein domain, the conservation of the amino acid, the Grantham score (26) and the designation of the mutation by three existing in-silico software tools; including a SIFT classification of “damaging” (27), and a Mutation Taster designation as “disease-causing” (28).

**Table 3.** Exome Variant Analysis.

| Familial Variant Analysis | # of Variants |
| --- | --- |
| Total Variants | 126,752 |
| Quality Control | 103,431 |
| Coding variants | 28,875 |
| Non-synonomous variants | 14,312 |
| European American variants with<br>MAF<0.01<br>(NHLBI ESP6500 exome data (20)) | 2,943 |
| Variants with MAF<0.01<br>(1000 genomes project (21)) | 1,801 |
| Variants with MAF<0.01<br>(CCHMC AA altering Variants (22)) | 769 |
| Recessive analysis | 1 |

The identified variant is a missense mutation in the gene *Coatomer Protein Complex, Subunit Beta 2* (*COPB2*). Consistent with the microarray data, this mutation was within the previously identified region of homozygosity on 3q22.1q24. We confirmed this exome result by Sanger Sequencing in all participants sequenced and extended the Sanger analysis to the unaffected sibling, who was heterozygous for the *COBP2* mutation (Fig 1K). The missense mutation in exon 8 of *COPB2* (c.760C>T; p.Arg254Cys) occurs at a highly conserved residue in a conserved region of the protein (Fig 1L). The affected amino acid is within a predicted WD40 protein-binding domain (Fig 2A). *Copb2* is highly expressed in the mouse ventricular zone (VZ) where neuroprogenitor cell division occurs in both mouse and human (29). This expression pattern is consistent with a model wherein reduction or loss of *COPB2* function could cause primary microcephaly.

**Figure 2.**
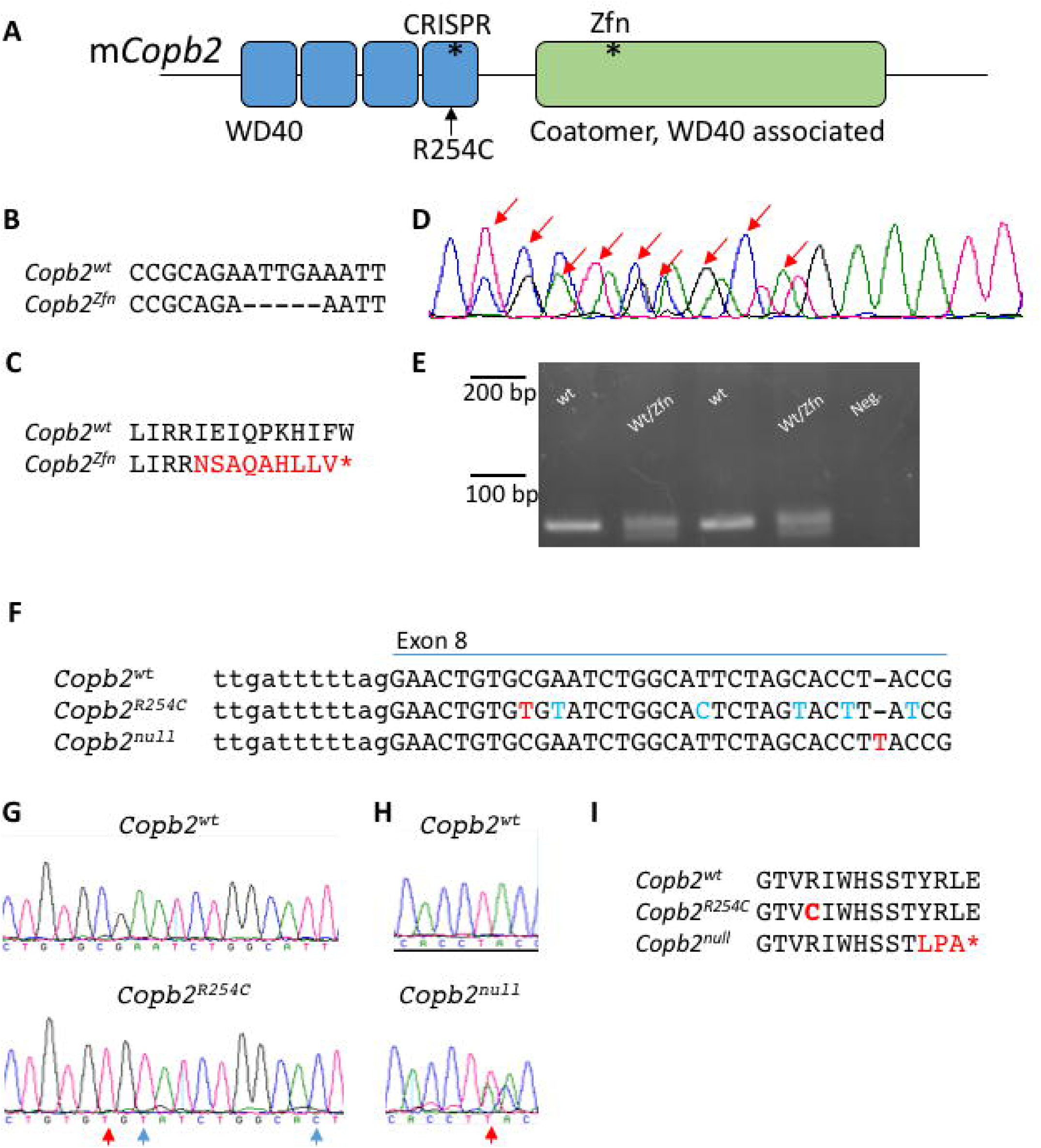
Mouse alleles of COPB2. (A) Schematic of the mouse COPB2 protein domains. Asterisks denote sites of mutations. (B-E) The *Copb2*^*Zfn*^ allele is a 5bp deletion in exon 12 (B,D), which results in a frameshift and creates a premature stop codon (C). (D) Sanger sequencing of the PCR products from a heterozygous animal. Sanger peaks that differ from wild-type are indicated by red arrows and this analysis identifies the precise nature of the deletion. (E) Genotyping was performed by PCR amplification of a 79bp region surrounding the deletion and gel electrophoresis (4% metaphor agarose). (F-H) The *Copb2^R^254^C^* (F,G) and *Copb2^null^* (F,H) alleles were created using CRISPR-Cas9 technology targeting exon 8. Red letters indicate nucleotide changes affecting the amino acid sequence while blue letters indicate silent mutations. (I) *Copb2^R^254^C^* creates an amino acid change orthologous to the patient mutation at amino acid position 254, while *Copb2*^*null*^ creates a frameshift resulting in a premature stop codon (*) at position 264.

### A Copb2 allelic series in mouse

In order to validate our results in a mammalian system and to further explore the role of *COPB2* in development, we generated several alleles in the mouse. The first allele, *Copb2^em^1^Rstot^* (*Copb2*^*Zfn*^), is a Zinc-Finger nuclease mediated 5bp deletion within exon 12 which encodes a premature stop codon ten amino acids downstream of the deletion (Fig 2A-D). PCR amplification of a 79bp region surrounding the deletion allowed us to distinguish the two species upon gel electrophoresis (Fig 2E). We then used CRISPR-Cas9 technology to generate a “humanized” allele, *Copb2*^*em*^2^*Rstot*^ (*Copb2^R^254^C^*), with the missense variant seen in the patients (Fig 2A,F,G,I). In the process of creating the *Copb2^R^254^C^* allele, animals were also created with a one base pair insertion downstream of desired missense mutation site, creating another premature stop, *Copb2*^*em*^3^*Rstot*^, (*Copb2*^*null*^; Fig 2A,F,H,I).

### Copb2 is indispensable for embryogenesis

Given the integral role *Copb2* plays in cellular trafficking as a subunit of the Golgi Coatomer Complex, we hypothesized that loss of *Copb2* would severely impair embryogenesis. Results from both of our presumed null alleles (*Copb*^*Zfn*^ and *Copb2*^*null*^) supported this hypothesis. Of 35 embryos from E11.5-E18.5, and 11 from E7.5-E10.5 recovered from *Copb2*^*Zfn/wt*^ intercrosses, no homozygous *Copb2*^*Zfn/Zfn*^ embryos were recovered. Similarly, no *Copb2*^*null/null*^ embryos were recovered at E12.5 (Table 4). In many cases, there was evidence of early embryonic resorption, consistent with early lethality.

**Table 4.**
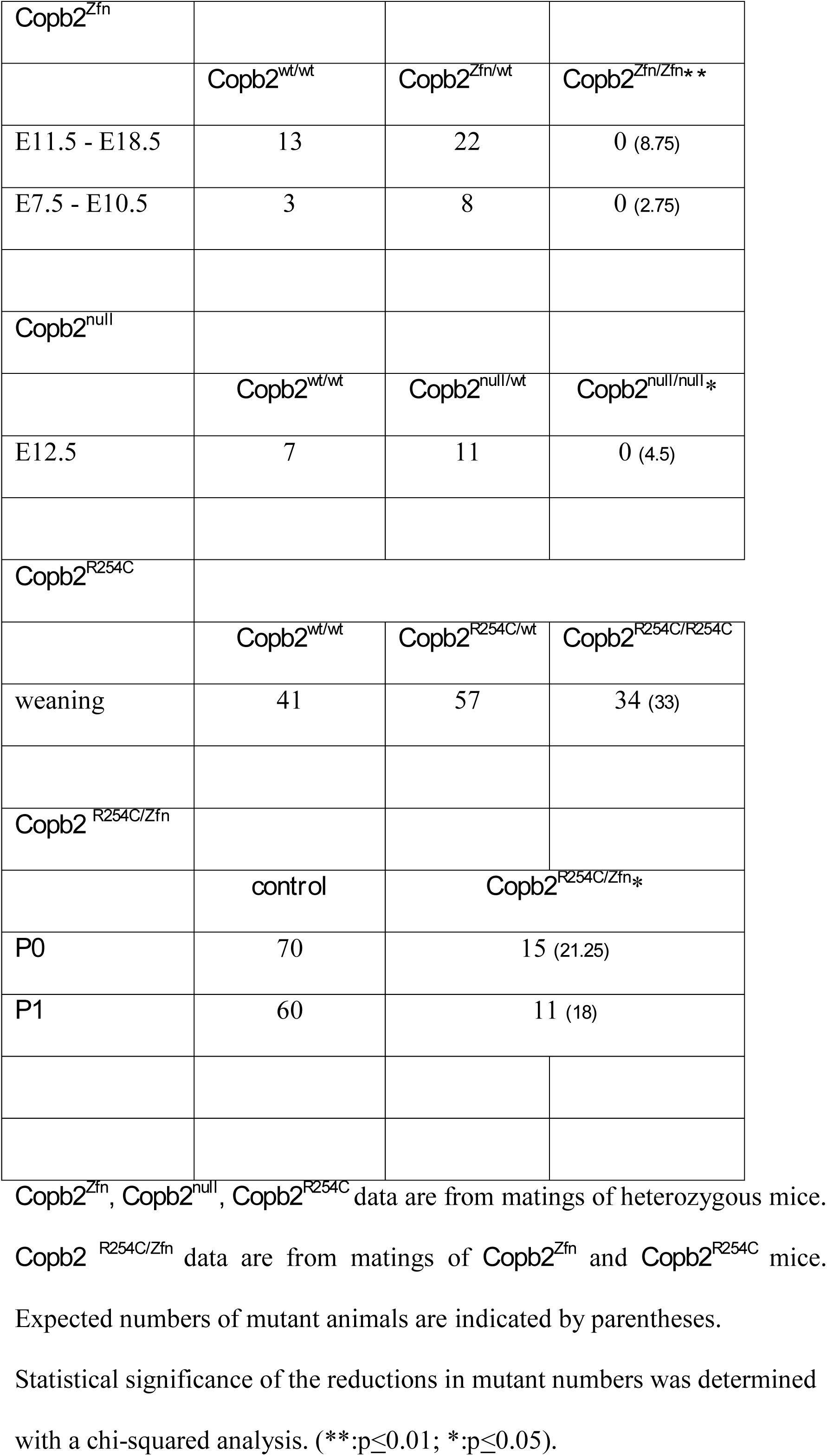
– Genotypes recovered from crosses of *Copb2* alleles.

|  |  |  |  |
| --- | --- | --- | --- |
| ***Copb2*<sup>Zfn</sup>** |  |  |  |
|  | *Copb2*<sup>wt/wt</sup> | *Copb2*<sup>Zfn/wt</sup> | *Copb2*<sup>Zfn/Zfn**</sup> |
| E11.5 - E18.5 | 13 | 22 | 0 (8.75) |
| E7.5 - E10.5 | 3 | 8 | 0 (2.75) |
| ***Copb2*<sup>null</sup>** |  |  |  |
|  | *Copb2*<sup>wt/wt</sup> | *Copb2*<sup>null/wt</sup> | *Copb2*<sup>null/null*</sup> |
| E12.5 | 7 | 11 | 0 (4.5) |
| ***Copb2*<sup>R254C</sup>** |  |  |  |
|  | *Copb2*<sup>wt/wt</sup> | *Copb2*<sup>R254C/wt</sup> | *Copb2*<sup>R254C/R254C</sup> |
| weaning | 41 | 57 | 34 (33) |
| ***Copb2*<sup>R254C/Zfn</sup>** |  |  |  |
|  | *control* | *Copb2*<sup>R254C/Zfn*</sup> |  |
| *P0* | 70 | 15 (21.25) |  |
| *P1* | 60 | 11 (18) |  |

*Copb2*<sup>Zfn</sup>, *Copb2*<sup>null</sup>, *Copb2*<sup>R254C</sup> data are from matings of heterozygous mice.
*Copb2*<sup>R254C/Zfn</sup> data are from matings of *Copb2*<sup>Zfn</sup> and *Copb2*<sup>R254C</sup> mice.
Expected numbers of mutant animals are indicated by parentheses.
Statistical significance of the reductions in mutant numbers was determined with a chi-squared analysis. (\*\*:p≤0.01; \*:p≤0.05).

### Copb2^R^254^C/R^254^C^ mice do not display any developmental abnormalities

Mice homozygous for the *R254C* allele are born in Mendelian ratios (Table 4) and are not physically distinct from littermate controls. Analysis of gross brain morphology and histology using Nissl staining did not reveal any discernable abnormalities in the brains of *R254C* homozygous or heterozygous animals compared to controls (Fig 3A-I). Body mass at both P21 and P60 was similar across all genotypes (Fig 3J). At 2 months of age, brain mass, forebrain area, and cell density appeared similar across all three genotypes (Fig 4K-M). Immunoblots for COPB2 show no significant decrease in protein levels in *Copb2^R^254^C/wt^* heterozygous or *Copb2^R^254^C/R^254^C^* homozygous animals (Fig 4N).

**Figure 3.**
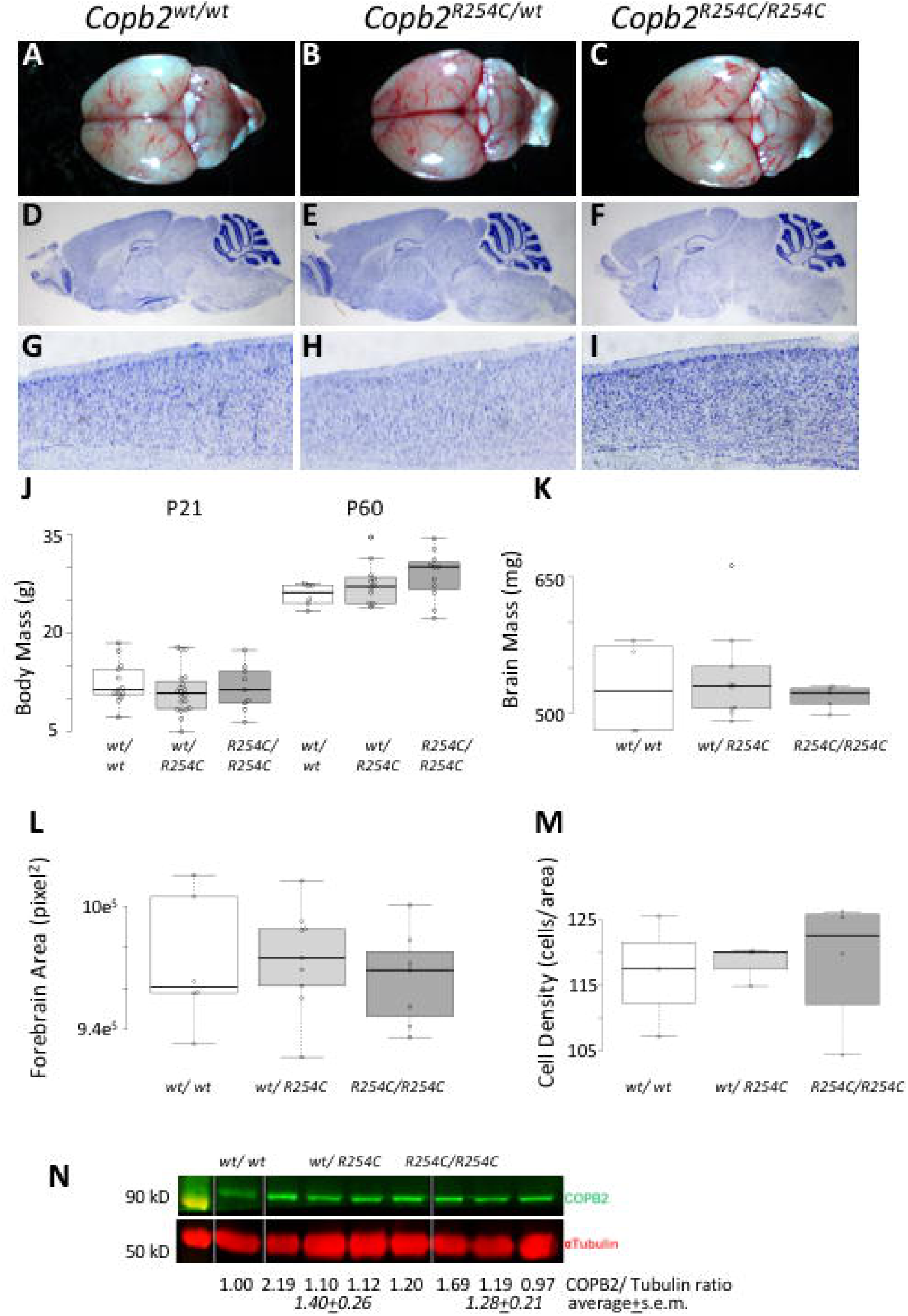
***COPB2R^254^C/R^254^C* mice are viable and do not have cortical malformations.** (A-C) Gross images of *Copb2^wt/wt^*(A), *Copb2^R^254^C/wt^*(B) *Copb2^R^254^C/R^254^C^*(C) brains at 2 months of age. (D-I) Histological Nissl stain shows similar brain structures across all three genotypes (J,K) Overall body and brain mass were similar between genotypes and forebrain area was not significantly different across genotypes (L). Additionally, high mag images of the cortex of all three genotypes (G-I) were quantified and showed no significant difference in cell density (M). For plots in J-M, center lines show the medians; box limits indicate the 25th and 75th percentiles as determined by R software; whiskers extend 1.5 times the interquartile range from the 25th and 75th percentiles, outliers are represented by dots; data points are plotted as open circles. Measurement of COPB2 protein showed similar levels across all three genotypes (N).

**Figure 4.**
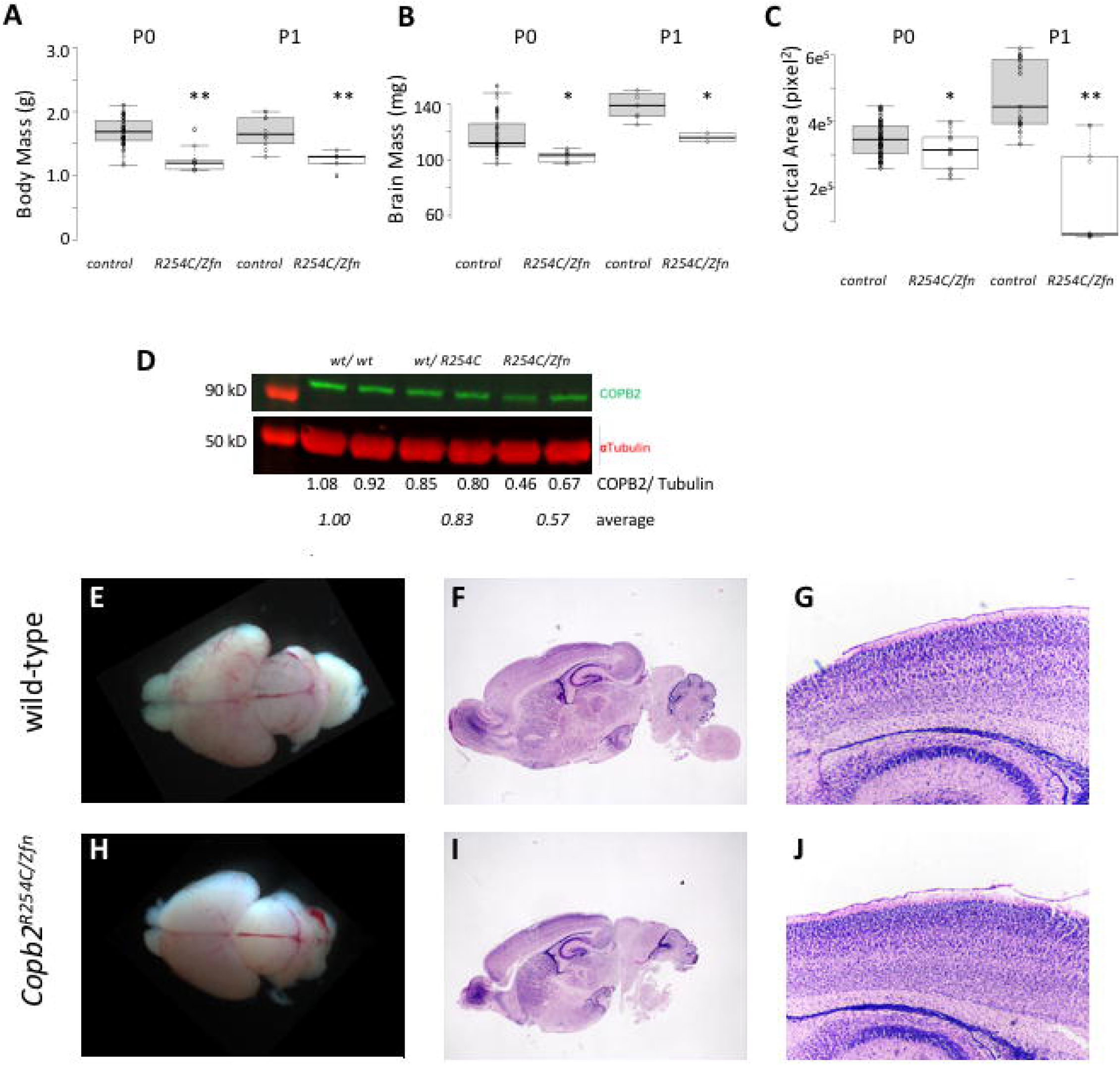
***COPB2R^254^C/Zfn* mice are perinatal lethal with reduced brain size and mass**. (A-B) Body and brain mass were significantly lower in *Copb2^R^254^C/Zfn^* mice compared to wild-type controls at both P0 and P1. (C) Cortical area was measured from gross images and showed a significant reduction in *Copb2^R^254^C/Zfn^* mice compared to control at both P0 and P1. For plots in A-C, center lines show the medians; box limits indicate the 25th and 75th percentiles as determined by R software; whiskers extend 1.5 times the interquartile range from the 25th and 75th percentiles, outliers are represented by dots; data points are plotted as open circles. Significance between groups was determined by a student’s t-test. (*:p<0.05; **:p<0.01). (D) COPB2 protein is reduced in *Copb2^wt/R^254^C^* and *Copb2^R^254^C/Zfn^* mice relative to wild-type controls. (E-J) Gross images of wild-type control (E) and *Copb2^R^254^C/Zfn^* brains (H) at P0. H&E stained sagittal sections of wild-type (F) and *Copb2^R^254^C/Zfn^* mice (I) mice show similar brain structure and similar cortical organization (G,J).

### Copb2^R^254^C/Zfn^ mice are perinatal lethal with cortical malformations.

As *Copb2^R^254^C/R^254^C^* mice appeared phenotypically normal, we hypothesized that the mouse brain is less sensitive to reductions in *Copb2* function than human. In order to test this hypothesis and further assess pathogenicity of the *R254C* allele, we generated *Copb2^R^254^C/Zfn^* animals. These pups were not born in Mendelian ratios (Table 4) and could be easily identified among littermate controls due to their small and sickly appearance, usually lacking a milk spot. Body mass was decreased in both P0 and P1 *Copb2^R^254^C/Zfn^* mice as compared to controls (Fig 4A). (With the exception of one litter in which all 4 *Copb2^R^254^C/Zfn^* survived the perinatal period but were morbid at P19, all *Copb2^R^254^C/Zfn^* animals were euthanized by P3 due to the excessive morbidity.) Additionally, both the mass and the cortical area of the *Copb2^R^254^C/Zfn^* brains were reduced compared to controls (Fig 4B, C). Western blots revealed a significant reduction in COPB2 protein levels in *Copb2^R^254^C/Zfn^* mice (approximately 50%; Fig 4D). Histological analysis of *Copb2^R^254^C/Zfn^* brains showed morphologically normal but reduced cortical tissues as compared to wild type controls (Fig 4E-J).

We began a molecular analysis of the *Copb2^R^254^C/Zfn^* phenotype to determine a potential mechanism for the reduction in cortical size. Immunohistochemistry for post-mitotic neurons using TuJ1 showed a robust population in both *Copb2^R^254^C/Zfn^* brains and controls (Fig 5A,B). Layer marker analysis was performed and we saw no reduction in TBR1-positive (layer VI) neurons in the *Copb2^R^254^C/Zfn^* brains compared to control (Fig. 5 C-E). In contrast, we saw a reduction in the later-born CTIP2-positive neurons (layer V) with a 32% reduction in *Copb2^R^254^C/Zfn^* perinatal brains (p<.05, Fig 5F-H). Immunostaining for apoptosis using Cleaved-Caspase 3 (CC3) antibodies at P0 showed significant increases in CC3-immunoreactivity in cortex of the *Copb2^R^254^C/Zfn^* animals (2.68 fold-increase relative to wild-type controls; Fig 5I-K). Increased cell death was also observed in the hippocampus, midbrain, cerebellum, and hindbrain of *Copb2^R^254^C/Zfn^* animals as compared to wild-type at P0-P3 (data not shown). *In vitro* experiments had indicated an increase in abortive autophagy following RNAi inhibition of *Copb2*, but we did not detect any change in the ratio of LC3-I to LC3-II levels in *R254C/Zfn* brains (Fig S1A).

**Figure 5.**
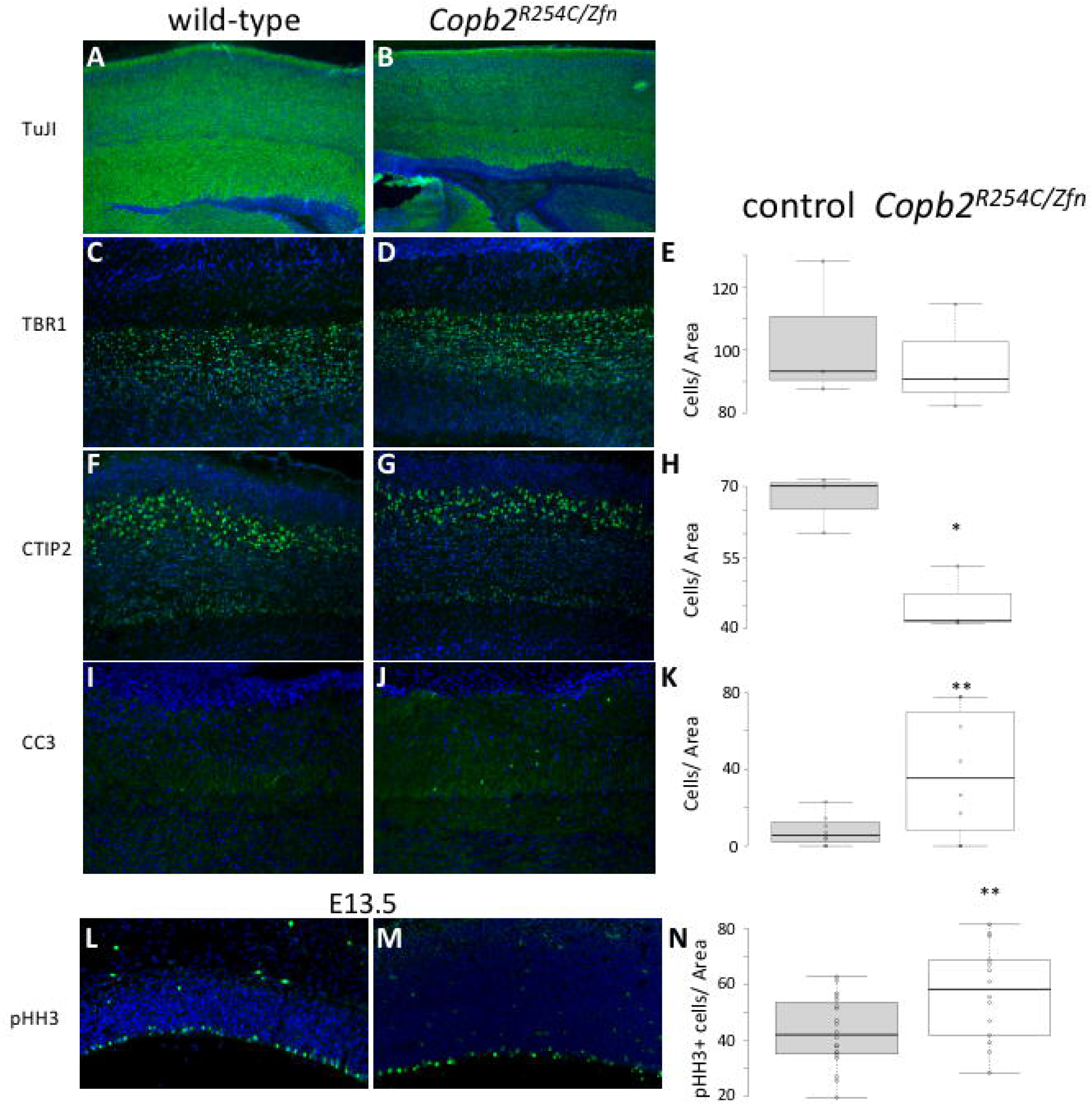
**Reduced layer V neuron production and increased cell death in *Copb2R*^254^*C/Zfn* brains**. (A-B) IHC for TuJ1-positive post-mitotic neurons highlights a robust population of cells in control (A) and *Copb2^R^254^C/Zfn^* mice (B). TBR1 IHC showed no significant reduction between control (C) and *Copb2^R^254^C/Zfn^* mice (D). CTIP2 IHC revealed a reduction in layer V neurons in *Copb2^R^254^C/Zfn^* mice (G) relative to control (F). Additionally, levels of CC3 to mark apoptotic cells were increased in the cortex of *Copb2^R^254^C/Zfn^* mice (J) compared to control (I). At E13.5, cell proliferation in the ventricular zone is increased in *Copb2^R^254^C/Zfn^* mice (L) compared to control (M). (E,H,K,N) Quantification of TBR1,CTIP2, CC3, and pHH3-immunoreactive cells was performed with an n = 3 for each genotype. Center lines show the medians; box limits indicate the 25th and 75th percentiles as determined by R software; whiskers extend 1.5 times the interquartile range from the 25th and 75th percentiles, outliers are represented by dots; data points are plotted as open circles. Significance between groups was determined by a student’s t-test. (*:p<0.05; **:p<0.01).

Finally, we began to interrogate neurogenesis at earlier developmental stages. We performed immunohistochemical analysis at E13.5 and E14.5 in the embryonic forebrain with phosphorylated histone H3 (pHH3) as a marker for proliferating cells. Quantification of pHH3 positive cells in the neurogenic ventricular zone revealed a slight but significant increase in *Copb2^R^254^C/ZnF^* proliferation as compared to littermate controls (33% increase, p<0.05, Fig 5L-N).

### Knockdown of Copb2 results in reduced proliferation in mouse neurospheres

Based on the decreased patient occipital head circumference and reduced brain size in the *Copb2^R^254^C/Zfn^* mice, we hypothesized that loss or reduction of COPB2 is associated with neural cell proliferation defects. Neurospheres were dissected from mouse embryos on embryonic days (E) 12.5-13.5 and cultured in DMEM-F12 medium with B27 and growth factors as described previously (30). We then used RNAi to deplete *COPB2* from NPCs and measured their proliferation. Over a three-day time course, control-infected NPCs increased in cell number over three-fold while *Copb2* silenced NPCs grew approximately 1.6-fold (Fig S2A). We also made neurospheres from the *Copb2* targeted animals and compared their ability to grow *in vitro.* In comparison to neurospheres from wild-type embryos, we saw reduced cell numbers in all genotypes tested at 3 days *in vitro* (Fig. S2B). At 5 days of culture, neurospheres from all *Copb2* genotypes still had fewer cells than the wild-type control. However, we noticed that neurospheres from *Copb2^R^254^C/Zfn^ mice* had significantly fewer cells than *Copb2^R^254^C/wt^* neurospheres, while the *Copb2*^*Zfn/wt*^ neurospheres had an intermediate number of cells. These data are consistent with our conclusions from the *in vivo* models that decreasing levels of *Copb2* are increasingly disrupting neural development.

To further confirm specific functional effects on *Copb2* of our RNAi constructs, we assayed for abortive autophagy, a known *Copb2* knockdown phenotype. We transfected nontargeted or *Copb2*-silenced intramedullary collecting duct (IMCD) cells with an LC3-RFP plasmid to measure autophagic flux. We used RFP-positive puncta in LC3-RFP transfected cells as a marker of autophagosome production, and GFP-RNAi constructs to identify cells with *Copb2* knockdown. In control IMCD cells, very few cells showed intense accumulation of LC3-RFP indicative of autophagic processing (Fig S3A). As a positive control, treatment with Brefeldin-A caused a significant accumulation of autophagosomes (Fig S3B). Cells treated with brefeldin-A (Fig S3B) or stably expressing the *Copb2* siRNA (as indicated by GFP, data not shown) showed significant increases in LC3-RFP puncta, indicative of increased numbers of autophagosomes (Fig S3C). We further labeled the cells with LysoTracker to mark the lysosomes. We first noted a significant increase in the number of lysosomes in *Copb2* knockdown cells as compared to controls (Fig. S3H). We also noted a significant amount of co-localization of the Lysotracker and the LC3-RFP, indicating fusion of the lysosomes and autophagosomes (Fig. S3I), consistent with previous results (31, 32). Taken together and in combination with previously published reports on the function of COPB2, these data show that reduction of COPB2 function leads to; 1) an increase in formation of autophagosomes which are not able to fully mature, 2) abortive autophagy and, 3) lowered cellular proliferation in mouse neurospheres. We were unable to generate clonal cell lines with either RNAi or CRISPR/CAS9 editing that lacked COPB2 entirely, consistent with our failure to recover homozygous null mouse embryos (Table 4) and previous literature (31).

## DISCUSSION

Here, we have identified a novel variant in the coding region of *COPB2* in two patients with primary microcephaly, seizures, and failure to thrive. To potentially demonstrate a role for *COPB2* in neural development and pathogenicity for this variant, we generated an allelic series in the mouse. We show that *Copb2* is required for survival to organogenesis stages. Strikingly, *Copb2^R^254^C/R^254^C^* mice appear phenotypically indistinct from littermates and are viable and fertile. However, reducing the proportion of functional *Copb2* further by generating *Copb2^R^254^C/Zfn^* mice leads to perinatal death and cortical malformations. In contrast to previous reports studying depletion of COPI components, the *Copb2^R^254^C/Zfn^* neural phenotype does not appear to involve an increase in abortive autophagy or ER stress. This study represents the first discovery of a disease-causing variant in *COPB2* as well as the first known link between primary microcephaly and a subunit of the COPI complex.

Like most genes linked to microcephaly, *COPB2* apparently acts as part of a highly conserved pathway that is essential for life in a wide variety of cellular lineages. This conclusion is supported by our discovery that *Copb2*^*null/null*^ and *Copb2*^*Zfn/Zfn*^ embryos do not survive to organogenesis (Table 4), and that our attempts to create COPB2-null cell lines were unsuccessful (data not shown). These results are also supported by the literature, where it has been demonstrated that siRNA-mediated reduction of *COPB2* in breast cancer cells leads to a drastic reduction in replating efficiency (31). Notably, an early lethality phenotype has also been observed in mice null for other known microcephaly-associated genes, including *MCPH7* (*STIL*), which die *in utero* after E10.5 (33), and *MCPH10 (ZNF335)*, which die as early as E7.5 (34). Thus, it appears human malformations are more likely due to hypomorphic mutations in these genes where neural development is especially susceptible, likely due to the extremely high rates of neurogenesis in primate cortical development.

The early lethality of *Copb2*^*null/null*^ embryos strongly suggests that the patient mutation is hypomorphic, as an effectively null allele would presumably arrest embryogenesis at a much earlier stage. This assumption proved true in our mouse models: immunoblots of perinatal *Copb2^R^254^C/R^254^C^* brains indicate no significant decrease in COPB2 levels, whereas *Copb2^R^254^C/ZnF^* brains have approximately half the normal amount of protein. The *COPB2^R^254^C^* mutation is located within the N-terminal WD40 repeat (β-propeller) domain of β’-COP, which has been shown to bind COPI cargo (35). Given the hypomorphic nature of the mutation, one could hypothesize that it interferes with the interaction between β’-COP and this cargo, causing an incremental reduction in COPI trafficking efficiency which is sufficient to cause defects during neurogenesis. Interestingly, the α-COP subunit of COPI shares significant N-terminal homology with β’-COP and has also been shown to bind cargo using its own WD40 repeat domain (36). *COPA* mutations in this domain result in Copa Syndrome, an immune regulatory syndrome in which the mutated α-COP protein has a reduced ability to bind di-lysine tagged cargo (37). It remains to be seen why *COPA* mutations result in such a syndrome while *COPB2* mutations in the homologous domain result in neurogenesis defects. Presumably, this can be attributed either to the fact that COPB2 binds a unique set of COPI cargo, or to the fact that the *COPB2^R^254^C^* mutation affects protein function to a different degree than known *COPA* mutations do. Future experiments to address differentially affected binding partners will be quite interesting in this regard.

Other hypotheses about the underlying mechanism could be based on studies of two other proteins harboring WD40-repeat domains. The first, WDR62, has reported mutations causing microcephaly due to mitotic delay and increased cell death (38). Additionally, another similar protein (WDR45) is associated with neurodegeneration, evidently through autophagy defects (39). However, the unchanged LC3-I/ LC3-II levels in *Cobp2*^*R*^254^*C/ZnF*^ brain lysates suggest that autophagy is relatively uninterrupted, suggesting this is an unlikely mechanism causing the phenotypes we report here. It is surprising we do not see evidence for deficits in autophagic flux as we and others have seen upon RNAi mediated depletion of *Copb2 in vitro* (31). We hypothesize that the autophagy defects seen in the knockdown paradigms may represent a more significant decrease in levels of COPB2 than those generated in the genetic models. Indeed, the homozygous null embryos may have autophagy phenotypes, but were lethal so early in development this cannot be tested. The role of *Copb2* in regulating autophagy and a possible mechanistic link to microcephaly remains an intriguing possibility for future investigation.

A major difference between the patients discussed here and the mouse models we created was the ability of the mice to tolerate the *Copb2^R^254^C/R^254^C^* genotype without an observable phenotype, only exhibiting major defects when overall *Copb2* expression was reduced by a *Copb2^R^254^C/Zfn^* genotype. This issue is not unique to our study, as microcephaly has been historically difficult to model in the mouse, perhaps because mouse corticogenesis only generates around 14 million neurons while the human cortex has around 16 billion (40). Additionally, neurogenesis in gyrencephalic species is much more reliant on basal progenitor proliferation (41), suggesting that phenotypes involving this cell type may not be easily recapitulated in lissencephalic models such as the mouse. In some mouse models of microcephaly, such as *MCPH1* (42), and *MCPH2* (*Wdr62*) (38), reductions in cortical size are accompanied by comparable reductions in other brain structures and in overall body size, as was observed in *Copb2^R^254^C/Zfn^* mice. Additionally, we did find a reduction in layer V CTIP2-positive neurons in the *Copb2^R^254^C/Zfn^* mice, which is a reported feature in mouse models of microcephaly (43). It is also worth noting that the probands themselves were significantly underweight in addition to being microcephalic (Fig 1).

In light of the reduction in neurons expressing the cortical layer V marker CTIP2 (Fig 5F-H) and the significant increase in CNS apoptosis in neonates (Fig 5I-K), the increased VZ proliferation in E13.5 *Copb2^R^254^C/ZnF^* brains is intriguing. Although a complete elucidation of mechanism is beyond the scope of this study, these results invite further investigation. Notably, E13.5 is the peak day for layer V neurogenesis (44). One potential explanation for an increase in VZ proliferation associated with a decrease in layer V is a discrepancy in the ratio of symmetrical to asymmetrical divisions among the progenitor cells (45). This hypothesis could be further tested in the future along with a comprehensive study of the neurodevelopment in these mutants. It also remains to be seen whether this prenatal pHH3 increase is accompanied by an increased apoptotic index before birth as has been seen in multiple models (e.g., (46)).

Although we found differences in proliferation rates in the genetically modified neurospheres (Fig S3B), these were not as striking differences between differing mouse genotypes or the deficits in the RNAi treated neurospheres. This could be due to inherent differences in neurospheres *in vitro* as compared to *in vivo* neurogenesis. It is notable that all *Copb2* allelic combinations show reduced neurosphere growth as compared to wild-type. At 5DIV, the differing genotypes are starting to diverge with the *Copb2^R^254^C/ZnF^* neurospheres showing a further significant decrease in cell number from *Copb2^R^254^C/wt^*, a heterozygote for a hypomorphic missense mutation. The reduced growth rate in *Copb2^R^254^C/ZnF^* neurospheres may decrease even further from the *Copb2*^*ZnF/wt*^ neurospheres (a heterozygote for a null mutation) if the cultures were maintained even further *in vitro*. Each of these genetically modified neurospheres may have more residual protein that the RNAi treated clones which shows striking proliferation defects although this was not explicitly tested in our study.

*Copb2* variants have not been connected to any human disease prior to this study, although mutations in *Copb2* as well as *Copa* (α-COP) and *Copb* (β-COP) have been linked to notochord defects in Zebrafish (47). Notably, a protein associated with neurodegeneration, SCYL1, was shown to interact with COPI subunits and to regulate retrograde Golgi to ER traffic (48). Also of note, COPI function (specifically, the δ-COP subunit) was implicated in the trafficking and metabolism of amyloid precursor protein, suggesting a role in Alzheimer’s progression (49).

## MATERIALS AND METHODS

### Subjects

Informed consent was obtained according to Cincinnati Children’s Hospital Medical Center (CCHMC) institutional review board protocol # 2012-0203. Following consent, whole blood was collected on the parents, residual DNA on the affected individuals, and spit on the unaffected sibling.

### Microarray

Affected siblings had microarray analysis performed using the Illumina Infinium Assay (San Diego, CA) with the Illumina HD Human OMNI-quad BeadChip platform. The chip contains approximately 1,140,419 probes to assess copy number variation and regions of homozygosity.

### Genotyping

Library generation, exome enrichment, sequencing, alignment and variant detection were performed in the CCHMC Genetic Variation and Gene Discovery Core Facility (Cincinnati, OH). Briefly, sheared genomic DNA was enriched with NimbleGen EZ Exome V2 kit (Madison, WI). The exome library was sequenced using Illumina’s Hi Seq 2000 (San Diego, CA). Alignment and variant detection was performed using the Broad Institute’s web-based Genome Analysis Tookit (GATK) (19). All analyses were performed using Genome Reference Consortium Build 37.

### Variant Filtering and Pathogenicity Assessment

Quality control and data filtering were performed on VCF files in Golden Helix’s SNP and Variation Suite (Bozeman, MT). Non-synonymous coding variants were compared to three control databases, including NHLBI’s ESP6500 exome data (20), the 1000 genomes project (21), EXAC Browser (25)and an internal CCHMC control cohort (22). Remaining variants were subject to autosomal recessive analysis with emphasis on homozygous recessive variants found in the region of homozygosity identified by SNP microarray. The identified variant was compared to known disease genes in the OMIM (REF) and Human Gene Mutation (HGMD) (23) databases, and to reported variants in dbSNP (24) and the Exome Variant Server. The variant was also analyzed using Interactive Biosoftware’s Alamut v2.2 (San Diego, CA) to determine location of mutation within a protein domain, the conservation of the amino acid, the Grantham score (26) and the designation of the mutation by three existing in-silico software tools, SIFT (27), Polyphen (50) and Mutation Taster (28). *COPB2* mutations were validated by Sanger Sequencing (see Supplemental Table 1).

### COPB2 RNAi and cell proliferation

RNAi constructs were purchased from the Thermo-Fisher pGIPZ collection. Lentivirus was made using standard transfection protocols of HEK293t cells and used to infect mIMCD cells. Cells were selected for knockdown by both puromycin resistance and GFP expression. For all hairpins, clonal lines were also created by limiting dilution. Cell proliferation assays were performed using the Cell Titer-Blue Cell Viability Assay (Promega). 10,000 IMCD cells were plated for each clone and cell titer assays were performed in triplicate 6 and 48 hours after plating. NPC proliferation assays were performed as previously described (51). Statistical significance was calculated with Student’s t-test. All antibodies were purchased from Sigma (St. Louis, MO).

### Autophagy Assays

IMCD cells which had undergone clonal selection for pGIPZ-RNAi constructs were further transfected with pmRFP-LC3 (Addgene: (52)). Three days after transfection cells were fixed and examined. As a positive control, Brefeldin-A was added at a final concentration of 10ug/mL for 1 hour before fixation. LysoTracker Deep Red (Molecular Probes) was used to visualize lysosomes by addition of a 50nM stock 2 hours before fixation. Note that LysoTracker is pseudocolored green to facilitate visualization but was imaged with a 647nm light source.

### Allele Generation

Zinc finger nuclease guides were purchased from Sigma (St. Louis, MO) and submitted to the CCHMC Transgenic Core for blastocyst injection. Founder animals were screened with PCR primers (Table 5). CRISPR guides targeting the region of interest within exon 8 (ENMUSE00000230688) of m*Copb2* were designed using the MIT CRISPR design tool (crispr.mit.edu). Three potential guide RNA (gRNA) sequences were selected and ordered as complementary oligonucleotide pairs with BbsI overhangs (Table 5). These were ligated into the pSpCas9(BB)-2A-Puro (px459) vector and transfected into 3T3 using the Lipofectamine® 3000 transfection reagent (Thermo Fisher Scientific, Massachusetts). 48 hours after transfection, the cells were harvested and genomic DNA was purified before being subjected to the Surveyor® mutation detection kit in order to test cutting efficiency (Integrated DNA Technologies, Coralville, IA). As a control, cutting efficiencies of potential m*Copb2* guides were compared with that of a previously-published m*Tet2* gRNA (Integrated DNA Technologies, Iowa). A donor sequence was designed to introduce the desired missense mutation *COPB2^R^254^C^* modeling the patients’ mutation (Table 5). The donor and the plasmid encoding the most efficient m*Copb2* gRNA (Guide 3, Table 5) were sent to the CCHMC Transgenic Core for blastocyst injections. Potential founders were validated with Sanger Sequencing. pSpCas9(BB)-2A-Puro (PX459) V2.0 was a gift from Feng Zhang (Addgene plasmid # 62988). All transgenic injections were done into animals from a mixed C57BL/6;DBA/2 background. (C57BL/6 x DBA/2, 2nd generation). *Copb2*^*Zfn*^ mice were maintained on a C57BL/6J background and all CRISPR/CAS9 alleles were maintained on a CD-1 background.

### Animal husbandry

All animals were housed under and approved protocol and standard conditions. All euthanasia and subsequent embryo/organ harvests were preceded by Isoflurane sedation. Euthanasia was accomplished via dislocation of the cervical vertebrae. For embryo collections, noon of the day of vaginal plug detection was designated as E0.5.

### Sample measurements (body mass, brain mass)

Sacrificed animals were weighed immediately following euthanasia. Control body weights in Fig. 4 are a compilation of all unaffected genotypes. Analyses of each discrete genotype did not reveal any significant differences between groups. Brains were removed, weighed, and imaged using a Zeiss SteREO Discovery.V8 stereomicroscope. Relative area was measured using Axiovision 40 software (v.4.8.2.0). Magnification scale was the same for samples of the same age. Box plots were generated with BoxPlotR (http://shiny.chemgrid.org/boxplotr/).

### Western immunoblotting

Following euthanasia, P0-P3 brains were dissected, flash-frozen, and weighed. These samples were then homogenized in RIPA buffer and stored at -80°C. Homogenized samples were diluted in 50:1 Laemelli Buffer: 2-Mercaptoethanol and heated to 90°C for 5 minutes. Samples were then loaded into Novex™ WedgeWell 10-16% Tris-Glycine gels and electroporated at 130V for 100-130 minutes in Tris-Buffered Saline. Transfers were performed on ice at 35V for 90 minutes in Transfer Buffer (25 mM Tris, 190 mM Glycine, 20% Methanol). The membranes were then washed in TBS and blocking was performed at room temperature in 1:1 Odyssey® Buffer: PBS. Primary antibody incubation was performed at 4°C overnight in 1:1 Odyssey® Buffer: PBS. Membranes were then washed three times in TBS. Membranes were then incubated in IRDye® 680RD and 800CW at room temperature for 1 hour and visualized using an Odyssey® CLx Imaging System. Analysis was performed using Image Studio Lite V5.2. Relative band intensity was compared to α-Tubulin loading controls in order to quantify results.

### Histological and immunohistochemical analysis

For adult histology and immunohistochemistry (IHC) adult (2 months of age) littermate animals underwent transcardial perfusion using cold heparinized phosphate buffered saline (PBS) and 4% paraformalydehyde (PFA) solution. Brains were dissected and fixed for 72 hours in 4% PFA at room temperature followed by immersion in 70% ethanol (for histology). Samples were then paraffin embedded, sectioned at 5um, and processed through hematoxylin and eosin (H&E) staining. Sections were sealed using Cytoseal Mounting Medium (Thermo-Scientific).

For postnatal IHC analysis, P0-P3 pups were euthanized and brains dissected and fixed overnight in 4% paraformaldehyde. Following this, samples were immersed in 30% sucrose solution for 24-48 hours until sufficiently dehydrated. Brains were embedded in OCT compound (Tissue-Tek) and sectioned at a thickness of 10uM. For IHC sections were immersed in PBS followed by blocking in 5% normal goal serum in PBST for one hour and overnight incubation in the following primary antibodies: TuJ1 (1:500, Sigma), TBR1 (1:500, Abcam), CTIP2 (1:200, Abcam), CleavedCaspase3 (1:300, Cell Signaling), and GFAP (1:500, Abcam). Sections were washed three times in PBS and incubated at room temperature in AlexaFluor 488 goat anti-rabbit or goat anti-mouse secondary (1:500, Life Technologies) for 1 hour followed by incubation in DAPI (1:1000) for 15 minutes. Sections were then rinsed in PBS and sealed with ProLong Gold Antifade Reagent (Life Technologies). For postnatal histology sections were rinsed in PBS followed by processing through H&E and Nissl staining. Sections were sealed using Cytoseal Mounting Medium (Thermo-Scientific). Embryos were harvested and cryoembedded using standard procedure. IHC for pHH3 (1:500, SIGMA) was as detailed above. All IHC sections were imaged on Nikon C2 Confocal microscope. All paired images were taken at the same magnification.

### Quantification of cortical cell number

Images were imported into Image J (Schneider et al., 2012) and areas were established around the region of interest in the cortex. Thresholding was used to detect specific populations in each channel (DAPI and GFP) and quantified using the Analyze Particles function. All cell measurements are normalized to the areas recorded in each respective sample.

## Acknowledgements

Funding. This work was supported by the Cincinnati Children’s Research Foundation and the National Institutes of Health (R01NS085023 to R.W.S.)

## Conflict of Interest

The authors do not declare any conflicts.

## Abbreviations

CCHMC: Cincinnati Children’s Hospital Medical Center
CC3: Cleaved-Caspase 3
COPB2: *Coatomer Protein Complex, Subunit Beta 2*
CNV: copy number variation
ER: endoplasmic reticulum
ExAC: Exome Aggregation Consortium
HGMD: Human Gene Mutation
IMCD: intramedullary collecting duct
NPC: neural precursor cells
pHH3: phosphorylated histone H3
ROH: regions of homozygosity
RNAi RNA: interference
VZ: ventricular zone

CM= Cisterna Magna

SF-L= Left Sylvian Fissure

SF-R= Right Sylvian Fissure

AP= anterior-posterior

T= transverse

## REFERENCES

(1) Gilmore, E.C. and Walsh, C.A. (2013) Genetic causes of microcephaly and lessons for neuronal development. Wiley Interdiscip. Rev. Dev. Biol., 2, 461–478.

(2) Basel-Vanagaite, L. and Dobyns, W.B. (2010) Clinical and brain imaging heterogeneity of severe microcephaly. Pediatr. Neurol., 43, 7–16.

(3) Alkuraya, F.S., Cai, X., Emery, C., Mochida, G.H., Al-Dosari, M.S., Felie, J.M., Hill, R.S., Barry, B.J., Partlow, J.N., Gascon, G.G. et al. (2011) Human mutations in NDE1 cause extreme microcephaly with lissencephaly [corrected]. Am. J. Hum. Genet., 88, 536–547.

(4) Bilguvar, K., Ozturk, A.K., Louvi, A., Kwan, K.Y., Choi, M., Tatli, B., Yalnizoglu, D., Tuysuz, B., Caglayan, A.O., Gokben, S. et al. (2010) Whole-exome sequencing identifies recessive WDR62 mutations in severe brain malformations. Nature, 467, 207–210.

(5) Bond, J., Roberts, E., Mochida, G.H., Hampshire, D.J., Scott, S., Askham, J.M., Springell, K., Mahadevan, M., Crow, Y.J., Markham, A.F. et al. (2002) ASPM is a major determinant of cerebral cortical size. Nat. Genet., 32, 316–320.

(6) Bond, J., Roberts, E., Springell, K., Lizarraga, S.B., Scott, S., Higgins, J., Hampshire, D.J., Morrison, E.E., Leal, G.F., Silva, E.O. et al. (2005) A centrosomal mechanism involving CDK5RAP2 and CENPJ controls brain size. Nat. Genet., 37, 353–355.

(7) Hussain, M.S., Baig, S.M., Neumann, S., Nurnberg, G., Farooq, M., Ahmad, I., Alef, T., Hennies, H.C., Technau, M., Altmuller, J. et al. (2012) A truncating mutation of CEP135 causes primary microcephaly and disturbed centrosomal function. Am. J. Hum. Genet., 90, 871–878.

(8) Jackson, A.P., Eastwood, H., Bell, S.M., Adu, J., Toomes, C., Carr, I.M., Roberts, E., Hampshire, D.J., Crow, Y.J., Mighell, A.J. et al. (2002) Identification of microcephalin, a protein implicated in determining the size of the human brain. Am. J. Hum. Genet., 71, 136–142.

(9) Kumar, A., Girimaji, S.C., Duvvari, M.R. and Blanton, S.H. (2009) Mutations in STIL, encoding a pericentriolar and centrosomal protein, cause primary microcephaly. Am. J. Hum. Genet., 84, 286–290.

(10) Poirier, K., Lebrun, N., Broix, L., Tian, G., Saillour, Y., Boscheron, C., Parrini, E., Valence, S., Pierre, B.S., Oger, M. et al. (2013) Mutations in TUBG1, DYNC1H1, KIF5C and KIF2A cause malformations of cortical development and microcephaly. Nat. Genet., 45, 639–647.

(11) Shen, J., Gilmore, E.C., Marshall, C.A., Haddadin, M., Reynolds, J.J., Eyaid, W., Bodell, A., Barry, B., Gleason, D., Allen, K. et al. (2010) Mutations in PNKP cause microcephaly, seizures and defects in DNA repair. Nat. Genet., 42, 245–249.

(12) Hu, W.F., Chahrour, M.H. and Walsh, C.A. (2014) The diverse genetic landscape of neurodevelopmental disorders. Annu. Rev. Genomics. Hum. Genet., 15, 195–213.

(13) Letourneur, F., Gaynor, E.C., Hennecke, S., Demolliere, C., Duden, R., Emr, S.D., Riezman, H. and Cosson, P. (1994) Coatomer is essential for retrieval of dilysine-tagged proteins to the endoplasmic reticulum. Cell, 79, 1199–1207.

(14) Orci, L., Palmer, D.J., Ravazzola, M., Perrelet, A., Amherdt, M. and Rothman, J.E. (1993) Budding from Golgi membranes requires the coatomer complex of non-clathrin coat proteins. Nature, 362, 648–652.

(15) Styers, M.L., O’Connor, A.K., Grabski, R., Cormet-Boyaka, E. and Sztul, E. (2008) Depletion of beta-COP reveals a role for COP-I in compartmentalization of secretory compartments and in biosynthetic transport of caveolin-1. Am. J. Physiol. Cell Physiol., 294, C1485–1498.

(16) Waters, M.G., Serafini, T. and Rothman, J.E. (1991) 'Coatomer': a cytosolic protein complex containing subunits of non-clathrin-coated Golgi transport vesicles. Nature, 349, 248–251.

(17) Fiedler, K., Veit, M., Stamnes, M.A. and Rothman, J.E. (1996) Bimodal interaction of coatomer with the p24 family of putative cargo receptors. Science, 273, 1396–1399.

(18) Jackson, L.P., Lewis, M., Kent, H.M., Edeling, M.A., Evans, P.R., Duden, R. and Owen, D.J. (2012) Molecular basis for recognition of dilysine trafficking motifs by COPI. Dev. Cell, 23, 1255–1262.

(19) McKenna, A., Hanna, M., Banks, E., Sivachenko, A., Cibulskis, K., Kernytsky, A., Garimella, K., Altshuler, D., Gabriel, S., Daly, M. et al. (2010) The Genome Analysis Toolkit: a MapReduce framework for analyzing next-generation DNA sequencing data. Genome research, 20, 1297–1303.

(20) Fu, W., O’Connor, T.D., Jun, G., Kang, H.M., Abecasis, G., Leal, S.M., Gabriel, S., Rieder, M.J., Altshuler, D., Shendure, J. et al. (2013) Analysis of 6,515 exomes reveals the recent origin of most human protein-coding variants. Nature, 493, 216–220.

(21) Genomes Project, C., Abecasis, G.R., Altshuler, D., Auton, A., Brooks, L.D., Durbin, R.M., Gibbs, R.A., Hurles, M.E. and McVean, G.A. (2010) A map of human genome variation from population-scale sequencing. Nature, 467, 1061–1073.

(22) Patel, Z.H., Kottyan, L.C., Lazaro, S., Williams, M.S., Ledbetter, D.H., Tromp, H., Rupert, A., Kohram, M., Wagner, M., Husami, A. et al. (2014) The struggle to find reliable results in exome sequencing data: filtering out Mendelian errors. Front. Genet., 5, 16.

(23) Stenson, P.D., Mort, M., Ball, E.V., Shaw, K., Phillips, A. and Cooper, D.N. (2014) The Human Gene Mutation Database: building a comprehensive mutation repository for clinical and molecular genetics, diagnostic testing and personalized genomic medicine. Human genetics, 133, 1–9.

(24) Sherry, S.T., Ward, M.H., Kholodov, M., Baker, J., Phan, L., Smigielski, E.M. and Sirotkin, K. (2001) dbSNP: the NCBI database of genetic variation. Nucleic Acids Res., 29, 308–311.

(25) Karczewski, K.J., Weisburd, B., Thomas, B., Solomonson, M., Ruderfer, D.M., Kavanagh, D., Hamamsy, T., Lek, M., Samocha, K.E., Cummings, B.B. et al. (2017) The ExAC browser: displaying reference data information from over 60 000 exomes. Nucleic Acids Res, 45, D840–D845.

(26) Grantham, R. (1974) Amino acid difference formula to help explain protein evolution. Science, 185, 862–864.

(27) Li, M.X., Gui, H.S., Kwan, J.S., Bao, S.Y. and Sham, P.C. (2012) A comprehensive framework for prioritizing variants in exome sequencing studies of Mendelian diseases. Nucleic Acids Res., 40, e53.

(28) Schwarz, J.M., Rodelsperger, C., Schuelke, M. and Seelow, D. (2010) MutationTaster evaluates disease-causing potential of sequence alterations. Nature methods, 7, 575–576.

(29) Diez-Roux, G., Banfi, S., Sultan, M., Geffers, L., Anand, S., Rozado, D., Magen, A., Canidio, E., Pagani, M., Peluso, I. et al. (2011) A high-resolution anatomical atlas of the transcriptome in the mouse embryo. PLoS biology, 9, e1000582.

(30) Dasgupta, B. and Milbrandt, J. (2009) AMP-activated protein kinase phosphorylates retinoblastoma protein to control mammalian brain development. Dev. Cell, 16, 256–270.

(31) Claerhout, S., Dutta, B., Bossuyt, W., Zhang, F., Nguyen-Charles, C., Dennison, J.B., Yu, Q., Yu, S., Balazsi, G., Lu, Y. et al. (2012) Abortive autophagy induces endoplasmic reticulum stress and cell death in cancer cells. PLoS One, 7, e39400.

(32) Razi, M., Chan, E.Y. and Tooze, S.A. (2009) Early endosomes and endosomal coatomer are required for autophagy. The Journal of cell biology, 185, 305–321.

(33) Izraeli, S., Lowe, L.A., Bertness, V.L., Good, D.J., Dorward, D.W., Kirsch, I.R. and Kuehn, M.R. (1999) The SIL gene is required for mouse embryonic axial development and left-right specification. Nature, 399, 691–694.

(34) Yang, Y.J., Baltus, A.E., Mathew, R.S., Murphy, E.A., Evrony, G.D., Gonzalez, D.M., Wang, E.P., Marshall-Walker, C.A., Barry, B.J., Murn, J. et al. (2012) Microcephaly gene links trithorax and REST/NRSF to control neural stem cell proliferation and differentiation. Cell, 151, 1097–1112.

(35) Eugster, A., Frigerio, G., Dale, M. and Duden, R. (2004) The alpha- and beta'-COP WD40 domains mediate cargo-selective interactions with distinct di-lysine motifs. Mol. Biol. Cell, 15, 1011–1023.

(36) Jackson, L.P. (2014) Structure and mechanism of COPI vesicle biogenesis. Curr. Opin. Cell Biol., 29, 67–73.

(37) Vece, T.J., Watkin, L.B., Nicholas, S.K., Canter, D., Braun, M.C., Guillerman, R.P., Eldin, K.W., Bertolet, G., McKinley, S.D., de Guzman, M. et al. (2016) Copa Syndrome: a Novel Autosomal Dominant Immune Dysregulatory Disease. J. Clin. Immunol., 36, 377–387.

(38) Chen, J.F., Zhang, Y., Wilde, J., Hansen, K.C., Lai, F. and Niswander, L. (2014) Microcephaly disease gene Wdr62 regulates mitotic progression of embryonic neural stem cells and brain size. Nat. Commun., 5, 3885.

(39) Haack, T.B., Hogarth, P., Gregory, A., Prokisch, H. and Hayflick, S.J. (2013) BPAN: the only X-linked dominant NBIA disorder. Int. Rev. Neurobiol., 110, 85–90.

(40) Koch, C. and Reid, R.C. (2012) Neuroscience: Observatories of the mind. Nature, 483, 397–398.

(41) Florio, M., Albert, M., Taverna, E., Namba, T., Brandl, H., Lewitus, E., Haffner, C., Sykes, A., Wong, F.K., Peters, J. et al. (2015) Human-specific gene ARHGAP11B promotes basal progenitor amplification and neocortex expansion. Science, 347, 1465–1470.

(42) Trimborn, M., Ghani, M., Walther, D.J., Dopatka, M., Dutrannoy, V., Busche, A., Meyer, F., Nowak, S., Nowak, J., Zabel, C. et al. (2010) Establishment of a mouse model with misregulated chromosome condensation due to defective Mcph1 function. PLoS One, 5, e9242.

(43) Sgourdou, P., Mishra-Gorur, K., Saotome, I., Henagariu, O., Tuysuz, B., Campos, C., Ishigame, K., Giannikou, K., Quon, J.L., Sestan, N. et al. (2017) Disruptions in asymmetric centrosome inheritance and WDR62-Aurora kinase B interactions in primary microcephaly. Sci. Rep., 7, 43708.

(44) Finlay, B.L. and Darlington, R.B. (1995) Linked regularities in the development and evolution of mammalian brains. Science, 268, 1578–1584.

(45) Siegenthaler, J.A., Ashique, A.M., Zarbalis, K., Patterson, K.P., Hecht, J.H., Kane, M.A., Folias, A.E., Choe, Y., May, S.R., Kume, T. et al. (2009) Retinoic acid from the meninges regulates cortical neuron generation. Cell, 139, 597–609.

(46) Stottmann, R.W., Donlin, M., Hafner, A., Bernard, A., Sinclair, D.A. and Beier, D.R. (2013) A mutation in Tubb2b, a human polymicrogyria gene, leads to lethality and abnormal cortical development in the mouse. Hum. Mol. Genet., 22, 4053–4063.

(47) Coutinho, P., Parsons, M.J., Thomas, K.A., Hirst, E.M., Saude, L., Campos, I., Williams, P.H. and Stemple, D.L. (2004) Differential requirements for COPI transport during vertebrate early development. Dev. Cell, 7, 547–558.

(48) Burman, J.L., Bourbonniere, L., Philie, J., Stroh, T., Dejgaard, S.Y., Presley, J.F. and McPherson, P.S. (2008) Scyl1, mutated in a recessive form of spinocerebellar neurodegeneration, regulates COPI-mediated retrograde traffic. J. Biol. Chem, 283, 22774–22786.

(49) Bettayeb, K., Hooli, B.V., Parrado, A.R., Randolph, L., Varotsis, D., Aryal, S., Gresack, J., Tanzi, R.E., Greengard, P. and Flajolet, M. (2016) Relevance of the COPI complex for Alzheimer’s disease progression in vivo. Proc. Natl. Acad. Sci U S A, 113, 5418–5423.

(50) Adzhubei, I.A., Schmidt, S., Peshkin, L., Ramensky, V.E., Gerasimova, A., Bork, P., Kondrashov, A.S. and Sunyaev, S.R. (2010) A method and server for predicting damaging missense mutations. Nat. Methods, 7, 248–249.

(51) Liu, X., Chhipa, R.R., Nakano, I. and Dasgupta, B. (2014) The AMPK Inhibitor Compound C Is a Potent AMPK-Independent Antiglioma Agent. Molecular cancer therapeutics, 13, 596–605.

(52) Kimura, S., Noda, T. and Yoshimori, T. (2007) Dissection of the autophagosome maturation process by a novel reporter protein, tandem fluorescent-tagged LC3. Autophagy, 3, 452–460.

(53) Garrett, W.J., Kossoff, G. and Warren, P.S. (1980) Cerebral ventricular size in children: a two-dimensional ultrasonic study. Radiology, 136, 711–715.

